# Histone Acetyltransferase KAT2A Stabilises Pluripotency with Control of Transcriptional Heterogeneity

**DOI:** 10.1101/347476

**Authors:** Naomi Moris, Shlomit Edri, Denis Seyres, Rashmi Kulkarni, Ana Filipa Domingues, Tina Balayo, Mattia Frontini, Cristina Pina

## Abstract

Cell fate transitions in mammalian stem cell systems have often been associated with transcriptional heterogeneity, however existing data have failed to establish a functional or mechanistic link between the two phenomena. Experiments in unicellular organisms support the notion that transcriptional heterogeneity can be used to facilitate adaptability to environmental changes and have identified conserved chromatin-associated factors that modulate levels of transcriptional noise. Herein, we show destabilisation of pluripotency-associated gene regulatory networks through increased transcriptional heterogeneity of mouse embryonic stem cells in which paradigmatic histone acetyl-transferase, and candidate noise modulator, Kat2a (yeast orthologue Gcn5) has been inhibited. Functionally, network destabilisation associates with reduced pluripotency and accelerated mesendodermal differentiation, with increased probability of transitions into lineage commitment. Thus, we functionally link transcriptional heterogeneity to cell fate transitions through manipulation of the histone acetylation landscape of mouse embryonic stem cells and establish a general paradigm that could be exploited in other normal and malignant stem cell fate transitions.

## INTRODUCTION

Cellular states, and their associated transitions, are a function of the transcriptional programmes active in the cell, which in turn depend on the chromatin configuration of their respective genes. It has been observed in multiple mammalian differentiation and developmental systems, including in mouse embryonic stem (ES) cells, that the cellular states during transition events are heterogeneous at the transcriptional level [1–7]. Such heterogeneity is thought to reflect temporal variability in gene expression, within individual cells, in an uncoordinated manner [8]. When variability affects genes that are regulators or effectors of cell fates, the expression status of individual genes can endow cells with different probabilities of effecting a transition [9], eventually resulting in a proportion of cells acquiring the new fate; thus, heterogeneity in expression of these genes would vary the transition probability from a given state.

The homeobox gene *Nanog* is a paradigmatic pluripotency regulator that exhibits such variability in gene expression [10–12]. *Nanog* is strictly required for establishment of pluripotency, both in vitro and in the embryo [13], but is dispensable for its maintenance [11]. *Nanog* transcriptional reporters have been used to prospectively isolate cells on the basis of expression levels and, while there is some reversibility between Nanog high and low expression states, Nanog low cells have a higher probability of exiting self-renewal into differentiation [10, 12]. A role of Nanog down-regulation in the probabilistic exit from pluripotency is supported by experiments coupling reversible Nanog knockdown with single-cell transcriptomics showing that remodelling of pluripotency networks associated with *Nanog* loss can be transiently reversed [14].

The Nanog transcriptional reporters that are based on stable GFP (heterozygous TNGA cells) [11] exhibit a tri-modal distribution of high, mid and low GFP populations. While the high and low states represent the active and inactive transcriptional state of Nanog, respectively, the mid-Nanog (MN) population is likely to contain cells in which the Nanog promoter has been recently switched off, reversibly or irreversibly, causing the GFP levels to decay. This population is less apparent in destabilised fluorescent reporters such as the destabilised Venus reporter line, Nanog-VNP [15], confirming that intermediate levels of expression are not sustainable and resolve rapidly into high (HN) or low (LN) states. Therefore, in theory, the MN population should encompass all bona fide early transition events out of pluripotency and into lineage commitment. However, its transient nature makes it difficult to probe the molecular programmes of the state transition separate from protracted GFP expression, or confounding dissociation between reporter and endogenous Nanog expression [16].

Assessing the mechanistic basis of the transition out of pluripotency can be finely achieved through the use of Nanog reporter systems and it may shed light on a putative contribution of transcriptional heterogeneity to the probabilistic nature of cell state transitions. Dynamic changes in transcriptional activity, and the resulting changes in state-transition probabilities, are likely to be regulated, at least in part, at the level of histone lysine acetylation. In yeast, amplitude and frequency of transcriptional bursting [17] are regulated by distinct histone acetyl-transferase and deacetylase complexes which determine levels of H3K9 acetylation (H3K9ac) in the promoter and the body of the gene [18]. Promoter acetylation influences transcriptional variability or noise, as measured by coefficient of variation (CV=standard deviation/mean). Loss of the histone acetyl-transferase Gcn5 or its partner Sgf29 result in increased noise, while loss of components of the Rdp3s HDAC complex, which increase levels of H3K9ac, reduce gene expression CV.

In order to achieve a global reduction of H3K9ac in mouse ES cells, we chemically inhibited Gcn5 with the MB-3 compound [19], which we have recently validated to phenocopy loss of Gcn5 homologue KAT2A in mammalian cells [20]. While this resulted in minimal changes to average gene expression levels, it caused a significant enhancement of expression heterogeneity for a number of genes, including Nanog, with associated remodelling of gene regulatory networks (GRNs). These changes associated with functional destabilisation of pluripotency and accelerated differentiation, namely to mesendodermal (ME) lineages. Our results suggest that increased transcriptional heterogeneity may not only reflect, but indeed mechanistically promote, state transitions in mammalian cells.

## MATERIALS AND METHODS

### Cell culture

E14Tg2A, TNGA [11], Nanog-VNP [15], Sox1-GFP [21] and T-GFP [22] mouse ES cells were maintained or differentiated as described [23]. 2i medium used N2B27 (NDiff 227, StemCells) with 1μM PDO325901 (R&D) and 3μM Chiron (Cambridge SCI). Kat2a inhibition used 100 μM MB-3 (ab141255, AbCam) [19], or an equal volume of DMSO. ES cell colony-forming capacity was quantified using Alkaline Phosphatase detection kit (Sigma Aldrich) as per manufacturer’s instructions.

### Flow Cytometry

Cell sorting was done on MoFlo (Beckman Coulter), FACSAria III (BD) or Influx (BD) machines, and analysis performed on a BD LSR-Fortessa, with DAPI as a dead cell marker. Apoptosis analysis used Annexin-V antibody (a13202 Fluor-568, Invitrogen) and DAPI as per manufacturers’ instructions. Cell cycle analysis used Propidium Iodide staining of ethanol-fixed cells.

### Wash-off Experimental Protocol

TNGA cells in ESLIF were grown for 1 or 2 days with DMSO or MB-3 (50 and 100 μM), or exposed to control N2B27 pro-differentiation conditions. At the end of the treatment, cells were washed of the treatment, their GFP profile assessed, and cultured for up to 5 days in ESLIF with daily monitoring of distribution of GFP levels (Supplementary Fig. 3a). The composite GFP distribution obtained from randomly sampled cells from all time-points and conditions was best described by 3 Gaussian curves that correspond to the HN, MN and LN states [24] (Supplementary Fig. 3b), and every cell was assigned to one of these states using a probabilistic soft clustering approach (see Supplementary Data). Population proportions were calculated for each condition at each time point, and kinetic modelling employed to estimate rates of transition between states and general state reversibility (Fig. 3b and c).

### RNA sequencing (RNA-seq)

RNA extracted from TNGA cells treated in triplicate for 48h in ESLIF with DMSO or MB-3 (100 μM), was used for polyA library preparation and sequencing on an Illumina HiSeq4000 instrument. Details of alignment, quantification, differential gene expression and data analysis are described in Supplementary Data. Data deposited in GEO (GSE114797).

### Chromatin immunoprecipitation (ChIP) sequencing

Chromatin was prepared from TNGA cells treated in duplicate for 48h in ESLIF with DMSO or MB-3 (100 μM), and immunoprecipitated with an anti-H3K9ac antibody (ab4441, AbCam), as decribed [20]. Details of alignment, peak calling and identification and data analysis are described in detail in Supplementary Data. Data deposited in GEO (GSE114797).

### Single-cell quantitative RT-PCR

Single-cell RT-qPCR followed the Fluidigm Two-Step Single-Cell Gene Expression method on a 96.96 Dynamic Array IFC and the BioMark HD system, with EvaGreen Supermix [25] (primers in Supplementary Table 1) or Taqman Assays (Supplementary Table 2). Commercially-available spike-ins (Fluidigm, C1 RNA Standards Assay Kit, PN100-5582) controlled for technical variability. Further details of analysis are described in Supplementary Data.

## RESULTS

### Kat2a inhibition in mouse ES cells decreases H3K9ac and enhances heterogeneity of Nanog expression

We treated TNGA cells with MB-3 and observed a global reduction of H3K9ac both in number and height (Fig. 1a) of acetylation peaks. As expected for H3K9ac mark, the large majority of peaks were located in the vicinity of transcriptional start sites (TSS) (Fig. 1b), and the regions in which acetylation was lost as a result of MB-3 treatment were significantly enriched for Kat2a/Gcn5 occupancy as per the ENCODE significance tool (Fig.1c), supporting specificity of MB-3 activity. Myc/Max and E2F binding was also enriched at these locations, highlighting their well-described cooperation with Kat2a [26, 27]. The loci specifically affected by Kat2a inhibition (Supplementary Files 1-2) associated with metabolic functions, namely mitochondrial, as well as DNA and RNA metabolism (Supplementary File 3), suggesting an impact on general, rather than tissue-specific, functions and putative pervasive activities across multiple cell types. Interestingly, the overall reduction in H3K9ac did not translate into substantial changes in gene expression (Fig. 1d), as RNA-seq analysis of MB-3 vs. DMSO (vehicle)-treated TNGA cells revealed 599 differentially expressed genes, with only 3 at a fold change ≥2 (Supplementary File 4). Differentially expressed genes showed overlap with genes with differential H3K9ac and were significantly enriched in Kat2a binding targets (Fig. 1e), namely those recently described in mouse ES cells [26] (Fig. 1f). Given that the reduction in H3K9ac was not matched by significant changes in gene expression levels, we asked whether treatment might instead result in enhanced cell-to-cell heterogeneity of transcription, as described for Gcn5 in yeast.

**Figure 1.**
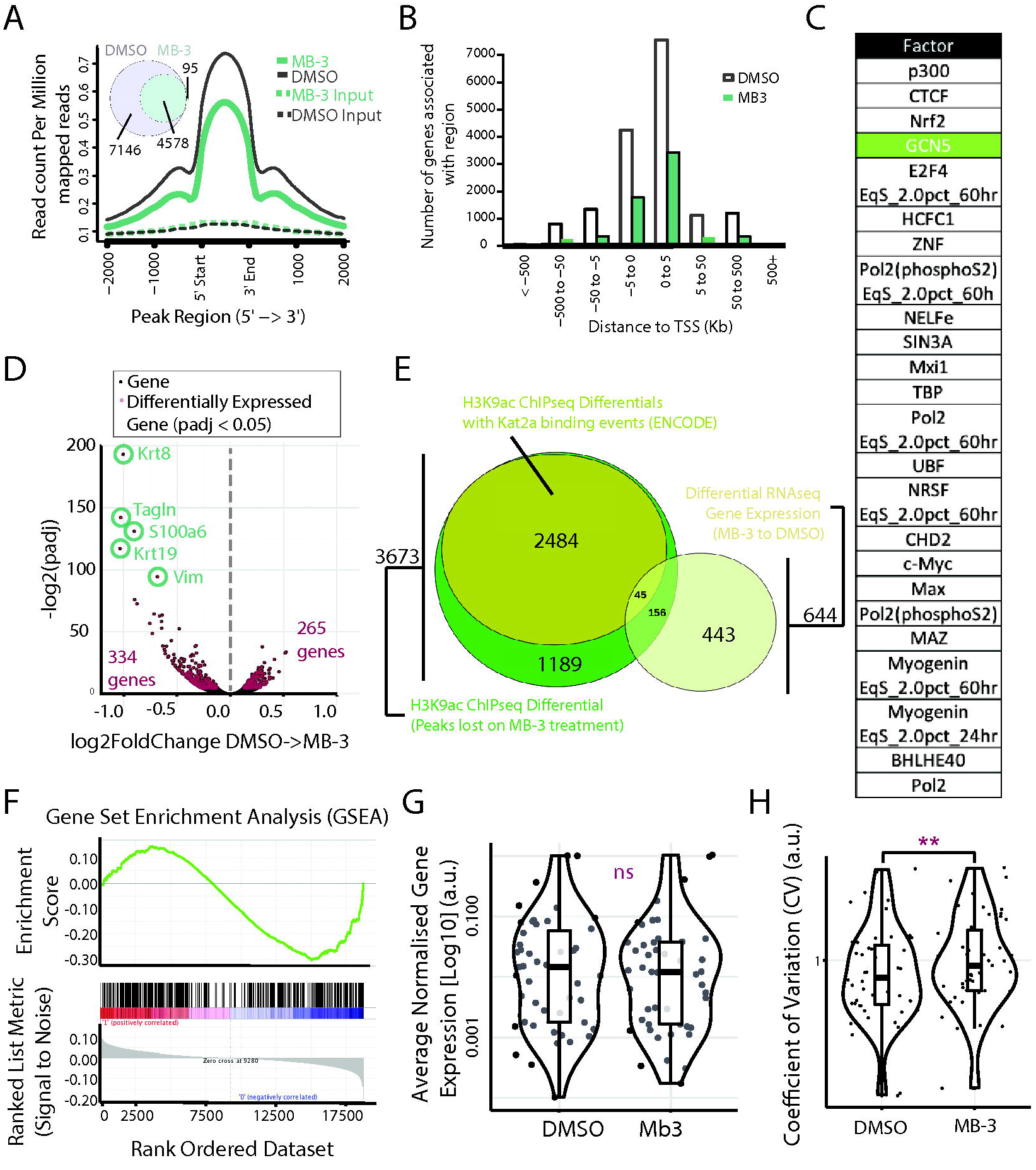
Effects of Kat2a inhibition on H3K9ac and transcriptional heterogeneity of mouse ES cells. **A**. MB-3 treatment of TNGA mouse ES cells results in specific loss of H3K9ac. Height and number (DMSO: 11724; MB-3: 4673, 98% overlap) of H3K9ac peaks are significantly reduced after 2 days of MB-3 treatment. **B**. H3K9ac marks associated with Transcriptional Start Sites (TSS) maintain their location profiles following MB-3 treatment but reduce in number. **C**. Transcription factor binding sites associated with genes that display reduced H3K9ac following MB-3 treatment include GCN5/Kat2A. List represents transcription factors with Q-value equal to 0, ordered by proportion of observed H3K9ac differential genes to total number of binding site-associated genes. **D**. RNA-seq analysis of TNGA mouse ES cells treated with MB-3 for 2 days. Volcano plot highlighting differentially-expressed genes (red dots: adjusted p-value < 0.05). **E**. Differentially expressed genes partially overlap with H3K9ac gene-associated peaks lost on MB-3 treatment. **F**. Gene set enrichment analysis of Kat2a direct targets in mouse embryonic stem cells amongst MB-3 down-regulated genes (ES = -0.30, NES = -1.34; p value = 0.005). **G**. Quantification of mean gene expression from scRT-qPCR from cells treated with DMSO or MB-3. There is no significant difference in mean gene expression between these conditions (Student’s t test, p>0.05). **H**. Coefficient of Variation (CV) of genes following scRT-qPCR shows increase in CV on MB-3 treatment (Student’s t test, p < 0.01).

We selected a representative subset of Kat2a binding targets, and interrogated 70 individual cells (35 DMSO, 35 MB-3) for transcript levels and cell-to-cell transcriptional variability (measured by coefficient of variation, CV), using single-cell quantitative RT-PCR (scRT-qPCR). Similar to the RNA-seq data, we did not find significant differences in mean expression levels between the 2 treatments (Fig. 1g). However, for the majority of the individual genes analysed, as well as at a global level, there was a significant increase in CV (Fig. 1h). The data thus supporting the hypothesis that inhibition of Kat2a-dependent H3K9ac results in increased transcriptional heterogeneity in mouse ES cells, similarly to previous observations in S. cerevisiae (supplementary data of [18]).

We then focused on components of the pluripotency network and asked whether their H3K9ac status was affected by Kat2a inhibition. We found that individual H3K9ac peaks were lost in the proximal upstream regions of the *Pou5f1/Oct4* locus [28] upon MB-3 treatment (Fig. 2a), an observation that extended to its regulator *Sall4* [29](Supplementary File 2). Furthermore, we observed a reduction in H3K9ac in the *Nanog* locus (Fig. 2a). The impact on Nanog prompted us to check whether Kat2a inhibition with MB-3 affected heterogeneity of GFP expression in TNGA cells, which has been associated with altered likelihood of pluripotency and differentiation state transitions [24]. Indeed, sustained inhibition of Kat2a activity modified GFP expression from the *Nanog* locus, with increased representation of MN cells and a broader distribution of fluorescence intensity values, distinct from the dominant HN observed in control cells (Fig. 2b). A similar increase in Nanog expression heterogeneity could be reproduced in wild-type E14tg2A mouse ES cells using immunofluorescence staining of endogenous NANOG protein (Fig. 2c), suggesting that the results were not limited to the TNGA genetic background.

**Figure 2.**
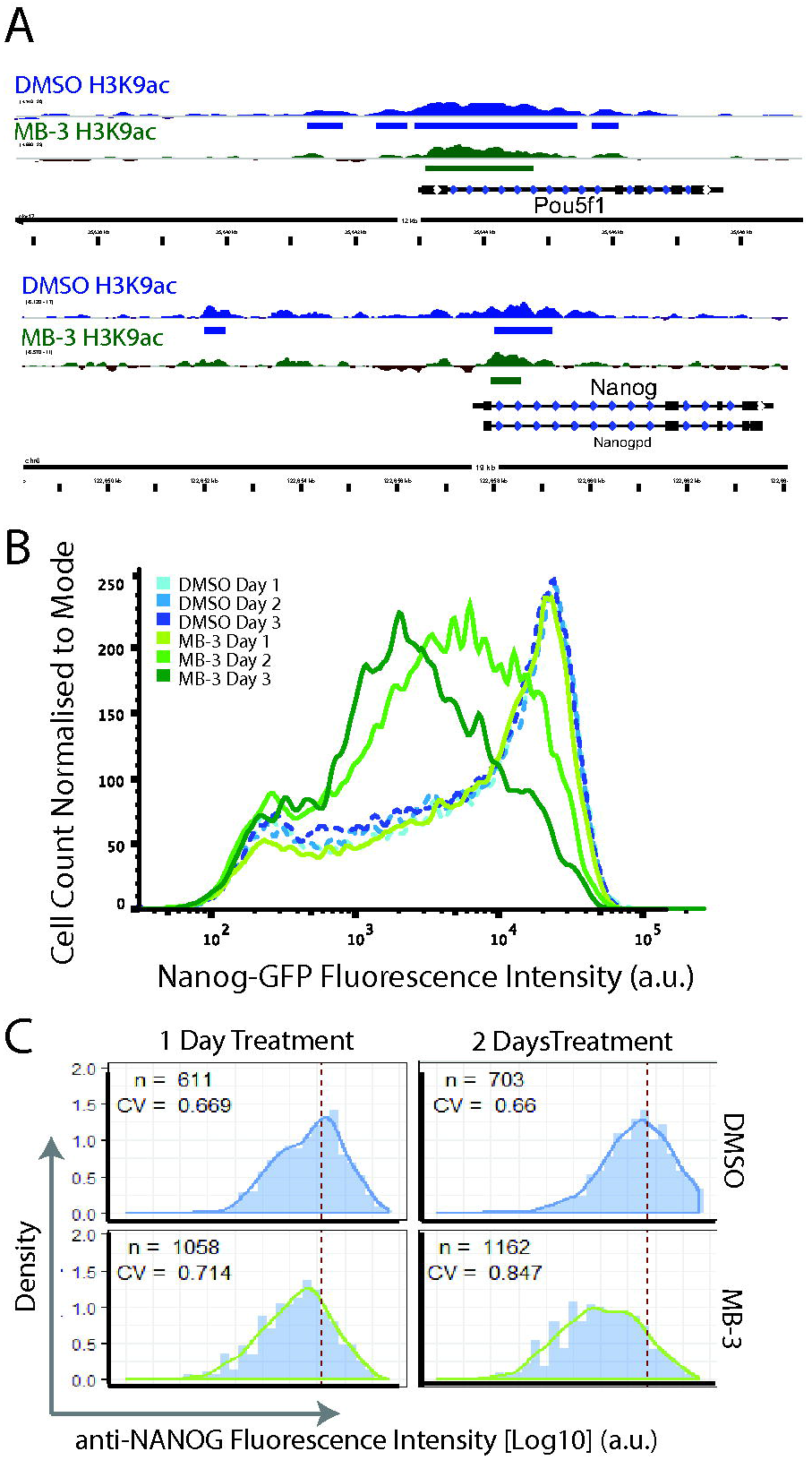
Effect of Kat2a inhibition on Nanog transcriptional heterogeneity. **A**. Chromatin immunoprecipitation (ChIP) analysis of H3K9 acetylation at the *Pou5f1* (top) and *Nanog* (bottom) loci, showing reduction in height and peak width/number upon MB-3 treatment. **B**. Flow cytometry analysis of heterozygous GFP expression from the *Nanog* locus in TNGA cells upon 1-3 days of MB-3 treatment; data representative >5 independent observations. **C**. Equivalent increase in heterogeneity observed in wildtype E14tg2A cells stained with NANOG antibody (see Supplementary Methods for quantification details).

Knockdown of Kat2a expression, through lentiviral-delivered shRNAs, supports the specificity of inhibitor activity. Kat2a depletion in TNGA cells increases heterogeneity in the distribution of Nanog levels in a manner proportional to the level of Kat2a knockdown, with accumulation of MN and LN cells (Supplementary Fig. 1a). Similar results are obtained in the Nanog-VNP cell line [15](Supplementary Fig. 1b), in which MB-3 treatment also results in a shift towards lower Nanog levels (Supplementary Fig. 1c). Overall, the results indicate that loss of Kat2a activity impacts variability of Nanog expression in a manner suggestive of enhanced transition out of pluripotency, as represented by high Nanog levels. We pursued these findings at a functional level to determine the associated impact on cellular state.

### Kat2a inhibition impacts mouse ES cell pluripotency and differentiation

We tested the functional impact of Kat2a inhibition on pluripotency by treating TNGA, E14tg2a and Nanog-VNP cells with MB-3 or DMSO in conventional serum and LIF-containing culture conditions (ESLIF; Fig. 3a). After 1-3 days of treatment, we washed off the inhibitor, and cultured the cells under stringent naïve pluripotency ‘2i’ conditions [30], after which we quantified the number of undifferentiated alkaline phosphatase-positive colonies obtained. While DMSO exposure had no effect on colony formation, cells exposed to MB-3 gradually lost the capacity to establish pluripotent colonies (Fig. 3b), compatibly with an increased probability of exiting the pluripotent state. In agreement with these findings, TNGA cells newly cultured in 2i conditions, in the presence of MB-3, show a broader GFP peak (Supplementary Fig. 2a), which is lost as cells undergo increased apoptosis (Supplementary Fig. 2b), presumably due to the inability of 2i medium to sustain primed and committed states. Cells cultured in ESLIF, on the other hand, do not show increased apoptosis (Supplementary Fig. 2b), and their cell cycle status is also not affected (Supplementary Fig. 2c). Interestingly, cells adapted to 2i conditions over several passages are unaffected by Kat2a inhibition (Supplementary Fig. 2d), which may reflect a change of epigenetic control, and is not dissimilar to loss of Nanog itself in mouse ES cells.

**Figure 3.**
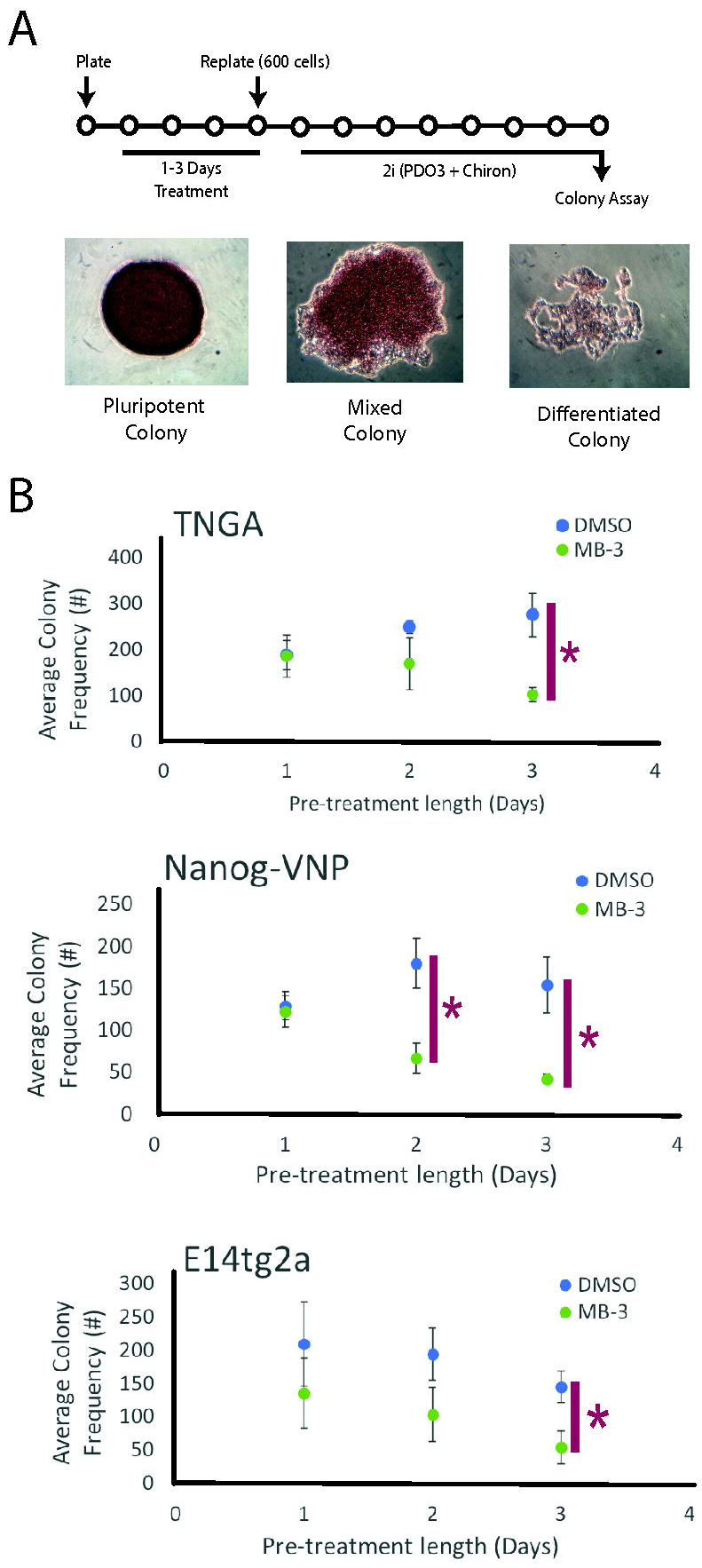
Effects of MB-3 treatment on naïve pluripotency. **A.** Experimental design: TNGA cells treated with MB-3 or DMSO in ESLIF (1-3 days), followed by passaging at clonal density and culture in 2i medium for 7 days. Pluripotency measured by colony number and alkaline phosphatase (AP) staining in colonies, as shown in representative images below. **B.** Significantly reduced number of AP+ pluripotent colonies from MB-3-treated cells after 2i medium switch (n=3; Student’s t test, p < 0.05). No significant difference was observed in frequencies of mixed or differentiated colonies (data not shown).

We then asked if Kat2a inhibition promoted differentiation decisions. We took advantage of ES cells with a GFP reporter of mesendodermal (ME) marker T (Brachyury) [22], and treated them for variable periods of time in the presence of DMSO or MB-3 before washing-off the treatment and placing the cells under ME-promoting culture conditions (Fig. 4a). We observed a significant acceleration of T expression in cells exposed to MB-3 (Fig. 4b). The same, however, was not true of Sox1-GFP [31] expression in ES cells placed under neuro-ectodermal (NE) culture conditions after inhibitor treatment (Supplementary Fig. 2e), suggesting some selectivity in lineage commitment decisions.

**Figure 4.**
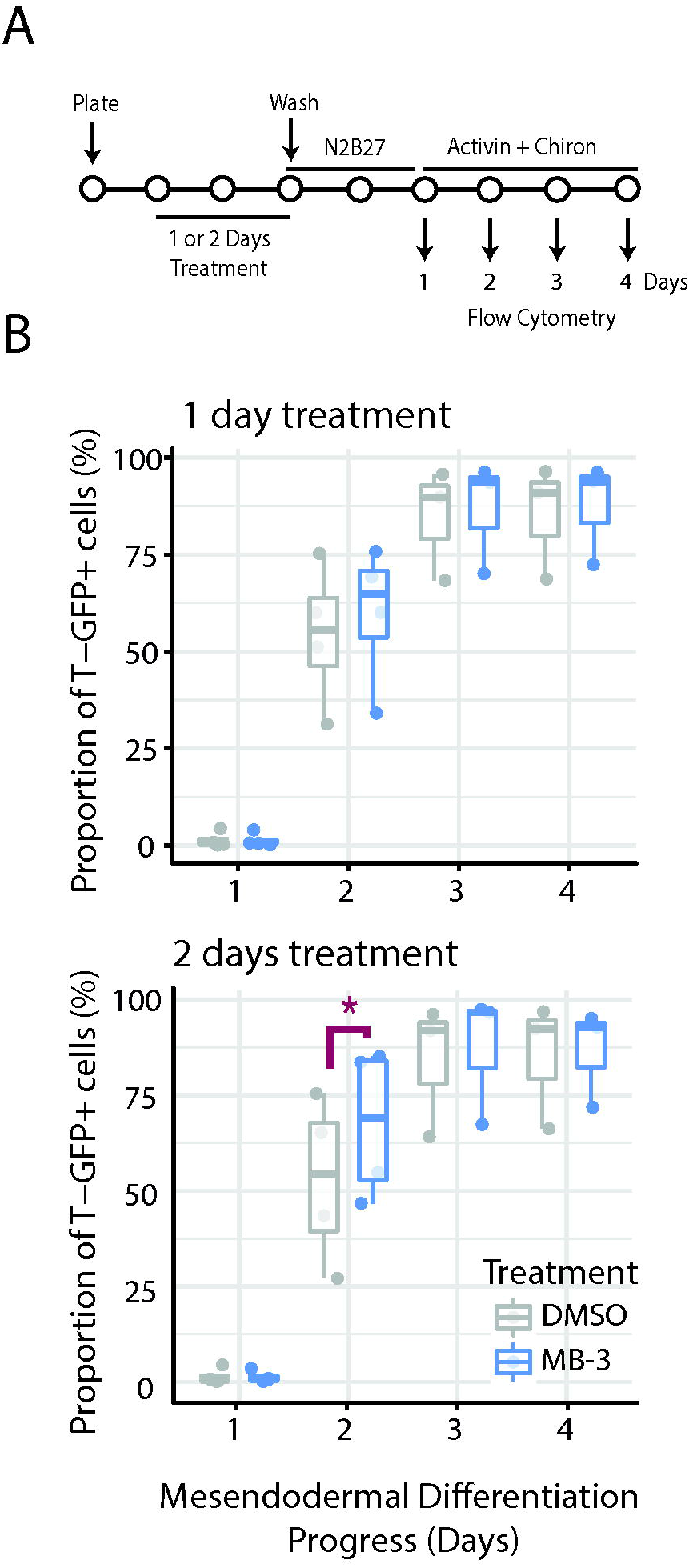
Effect of MB-3 treatment on mesendodermal differentiation of mouse ES cells. **A.** Mesendodermal Differentiation Experimental design: Brachyury-GFP (T-GFP) cells treated with MB-3 or DMSO in ESLIF medium (1-2 days) before wash-off and replating in N2B27 (Days 0-1) followed by N2B27 with 100ng/ml Activin and 3 μM Chiron (Days 2-4). Flow cytometry was used to quantify the proportion of GFP+ cells on Days 1-4 of differentiation. **B.** Exposure of mouse ES cells to MB-3 for 2 days significantly anticipates detection of T-GFP expression, indicative of accelerated ME commitment (n=4). Paired Student’s t-test *p<0.05

### Kat2a inhibition captures a mid-Nanog transition state in mouse ES cells

The impact of MB-3 treatment on pluripotency and ME differentiation is suggestive of a change in the probability of state transition, rather than with an absolute requirement to either state. To measure the probability of ES cells moving from pluripotency towards priming and commitment and the reversibility of these transitions, we used the TNGA cells and modelled rates of transition between HN (pluripotent), MN (primed), and LN (committed) states upon treatment exposure and wash-off (Fig. 5a; see Methods for details).

**Figure 5.**
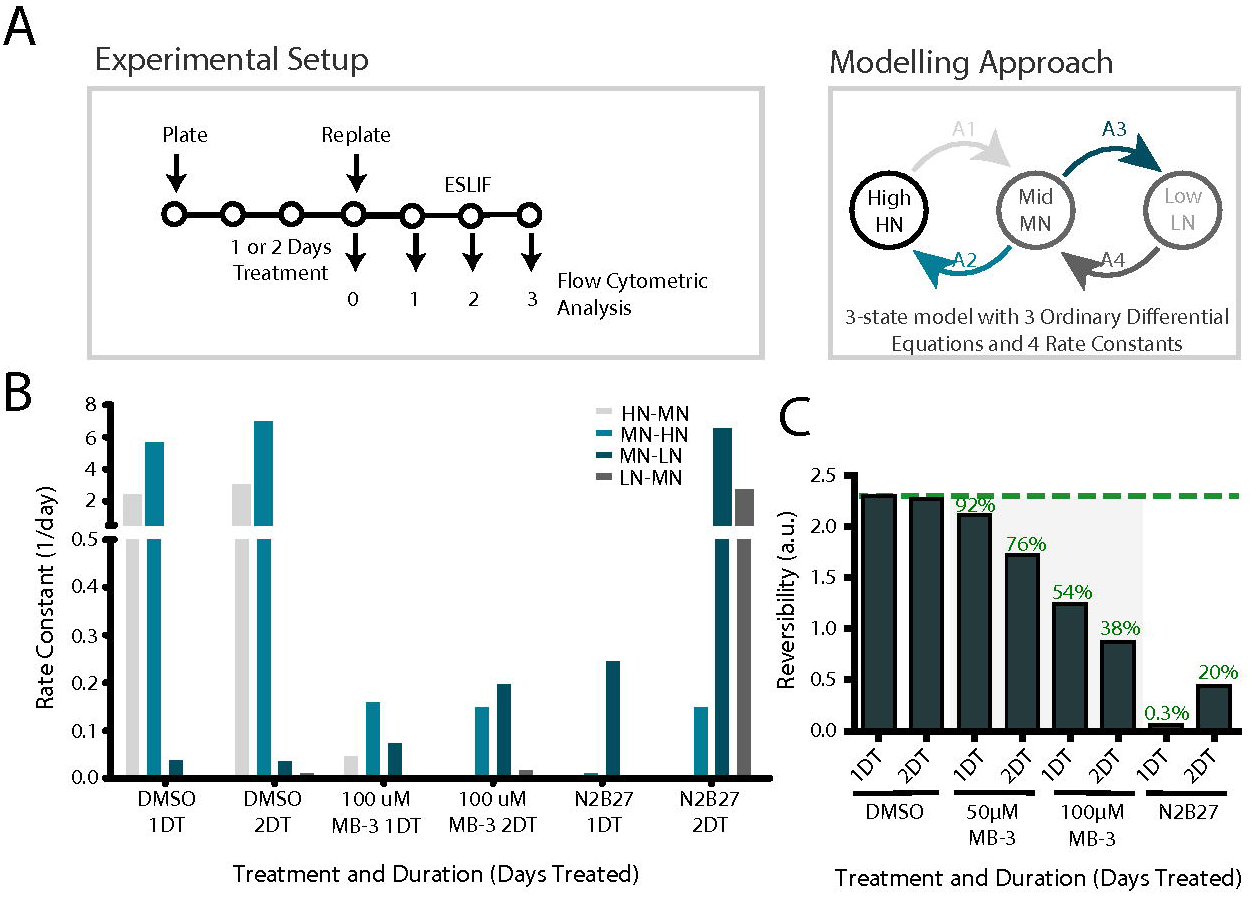
Inhibition of Kat2a catalytic activity affects reversibility of commitment decisions in TNGA mouse ES cells. **A**. Experimental design of wash-off experiments (left) and modelling approach (right). TNGA cells were cultured for 1-2 days in ESLIF medium with MB-3 (50 or 100 μM) or DMSO, or in N2B27 differentiation medium. Cells were washed off the treatment, and cultured for up to 3 days in ESLIF, with daily monitoring of Nanog-GFP profile. A model was constructed using the proportion of cells in each of the three High Nanog-GFP (HN), Mid Nanog-GFP (MN) and Low Nanog-GFP (LN) states with 4 kinetic parameters (A1-4; see Supplementary Methods). **B**. Kinetic modelling of transition rates between HN, MN and LN states in response to transient exposure to MB-3 (DMSO or N2B27, controls). HN-to-MN represents exit from pluripotency/commitment to differentiate; MN-to-LN represents differentiation; reversibility of either decision is given by the reverse transition probability. **C**. Reversibility indices (MN-to-HN + LN-to-MN) / (HN-to-MN + MN-to-LN) for different doses and durations of treatment (DT: Days Treatment). Green text indicates equivalent percentage of DMSO treatment.

DMSO-treated cells were unchanged by the treatment and transited between HN and MN states with a slight advantage towards the reverse movement, sustaining a dominant HN population. N2B27-treated cells showed a clear movement towards the LN state, denoting irreversible differentiation, which was dependent on the duration of treatment. MB-3 treated cells displayed a unique pattern of long-term retention in the MN state, with slow transition rates in either direction, and a progressive dose and time-dependent decrease in reversibility (Fig. 5b-c and Supplementary Fig. 3a-b). A similar pattern was observed in TNGA sorted as MN and exposed to MB-3 (Supplementary Fig. 3c). ◻ Overall the data suggest that inhibition of Kat2a-mediated acetylation permits the capture of mouse ES cells at the exit of pluripotency; in doing so, it may enrich for an otherwise fleeting state of transition out of pluripotency and into early ME differentiation.

### MB-3-driven pluripotency-to-ME transition associates with transcriptional heterogeneity

We next sought to characterise the transcriptional architecture of the transition state at singlecell resolution by interrogating individual cells captured from MB-3 and DMSO-treated cultures of mouse TNGA ES cells. We used scRT-qPCR transcriptional profiling and determined the presence, level, and heterogeneity of expression of genes central to ES cell pluripotency and differentiation (Supplementary Fig. 4a). We excluded LN cells to avoid the confounding effect of irreversibly committed or differentiated cells, and analysed 90 TNGA cells from ESLIF cultures, which are permissive to priming, and 138 from recently established naïve 2i cultures, which respond to MB-3 treatment with a broadened peak of GFP fluorescence intensity (Supplementary Fig. 2a), but do not sustain priming of differentiation programmes. ◻ Global representation of the transcriptional programmes of individual TNGA cells under the different experimental conditions shows an evident separation between the transcriptional spaces occupied by naïve and primed pluripotency. Importantly, it also reveals that the transcriptional space occupied by MB-3 treated 2i cells is in closer proximity to primed pluripotency than control (DMSO-treated) 2i cells (Fig. 6a), suggesting a requirement for Kat2a activity in establishing or maintaining pluripotency. In contrast, Kat2a active and inactive primed pluripotency transcriptional landscapes are closely overlapping (Fig. 6a), which may partly reflect the exclusion of cells with lower levels of Nanog, and also denote the slow transit of MB-3 treated cells out of the primed self-renewal (Supplementary Fig. 3b).

**Figure 6.**
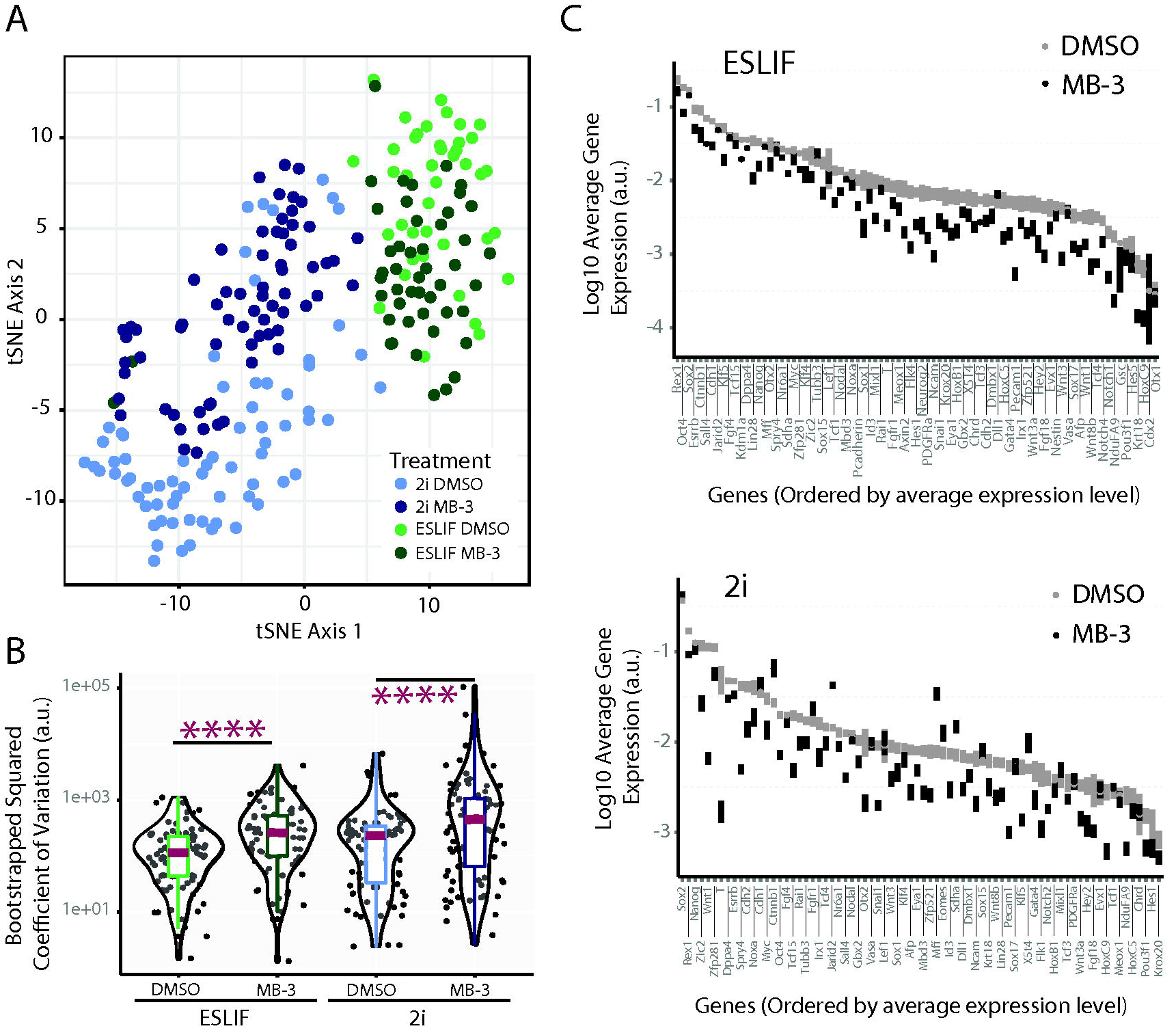
MB-3 treatment of TNGA mouse ES cells results in increased transcriptional heterogeneity from key pluripotency and differentiation regulatory loci. **A**. t-SNE plot of transcriptional profiles of individual TNGA cells treated with DMSO or MB-3 in ESLIF and 2i conditions. **B**. Overall changes in gene expression Squared Coefficient of Variation (CV2) between DMSO and MB-3-treated cells assayed from ESLIF (left) and 2i (right) culture conditions (Paired Student’s t-test p <0.0001). **C**. Ordered representation of gene expression changes between DMSO and MB-3-treated TNGA cells in ESLIF (top) and 2i (bottom) conditions. Mean expression represented as dot, standard error of the mean is represented as vertical bars.

We inspected the specific changes in single-cell gene expression patterns imposed by MB-3 treatment which underlay a misconfiguration of the 2i transcriptional state. We noted that MB-3 treated naïve pluripotent cells significantly enhanced cell-to-cell variability, or heterogeneity, of gene expression levels relative to control (Fig. 6b and Supplementary File 5), a finding reminiscent of the role of Kat2a yeast homologue Gcn5 in transcriptional noise [32]. A similar significant gain of gene expression variability was also observed in primed cells upon MB-3 treatment (Fig. 6b and Supplementary File 5). Increased heterogeneity occurred at all levels of average gene expression (Supplementary Fig. 4b), and with minimal changes to average gene expression levels (Fig. 6c), as observed in our bulk RNA-seq analysis. Frequency of gene expression was more dynamic between conditions, with minimal changes in ES-LIF, and some significant gains and losses in 2i in the presence of MB-3 (Supplementary Fig. 4c). We sought to articulate changes in gene expression frequency and variability of transcript levels through inference of gene regulatory networks (GRN), to understand the contribution of transcriptional heterogeneity to regulatory programmes at the exit of pluripotency.

### MB-3-driven reconfiguration of gene regulatory networks is centred on noise and H3K9ac changes

We focused on control and MB-3 treated cells in 2i culture conditions, where the transcriptional distance between states is more notable, and attempted to infer GRNs through a combination of binary and correlation methods we recently employed in an adult differentiation system [33] (Supplementary File 6). Given the proposed association from our data between loss of Kat2a-dependent H3K9ac and gene expression variability, we asked if this phenomenon might be instrumental in reconfiguring GRNs. Reassuringly, the inferred control network was strongly nucleated in pluripotency factor *Oct4/Pou5f1* (Fig. 7a), as well as in Wnt signalling elements *Ctnnb* and *Cdh1*, which likely reflect GSK-dependent Wnt activation in 2i culture conditions. Upon MB-3 treatment, we observed a specific reduction of network connectivity amongst nodes dependent on Kat2a activity for H3K9ac (Fig. 7b). Pluripotency regulators *Pou5f1* and *Sall4*, as well as *Jarid2* showed complete loss of connectivity upon Kat2a inhibition, whilst the network around *Cdh1* was greatly reduced (Fig. 7a-b). Importantly, when focusing on remodelled edges built around the respiratory chain component *Ndufa9* and differentiation factor *Mff*, both H3K9ac differential targets, we observed that the new connections established upon MB-3 treatment were with genes presenting significantly higher CV gains relative to non-remodelled edges (Fig. 7c); as a trend, gains in CV were also higher relative to edges exclusively detected in DMSO-treated cultures. A similar pattern of higher CV gains upon Kat2a inhibition was not observed in remodelled edges around nodes not strictly dependent on Kat2a for H3K9ac (Fig. 7d), suggesting a direct contribution to locus regulation. Furthermore, in interrogating the networks for genes exclusively connected upon Kat2a inhibition, we found that these presented higher CV gains than those genes exclusively present in the control network (Fig. 7e), suggesting a role for noise in network maintenance and reconfiguration.

**Figure 7.**
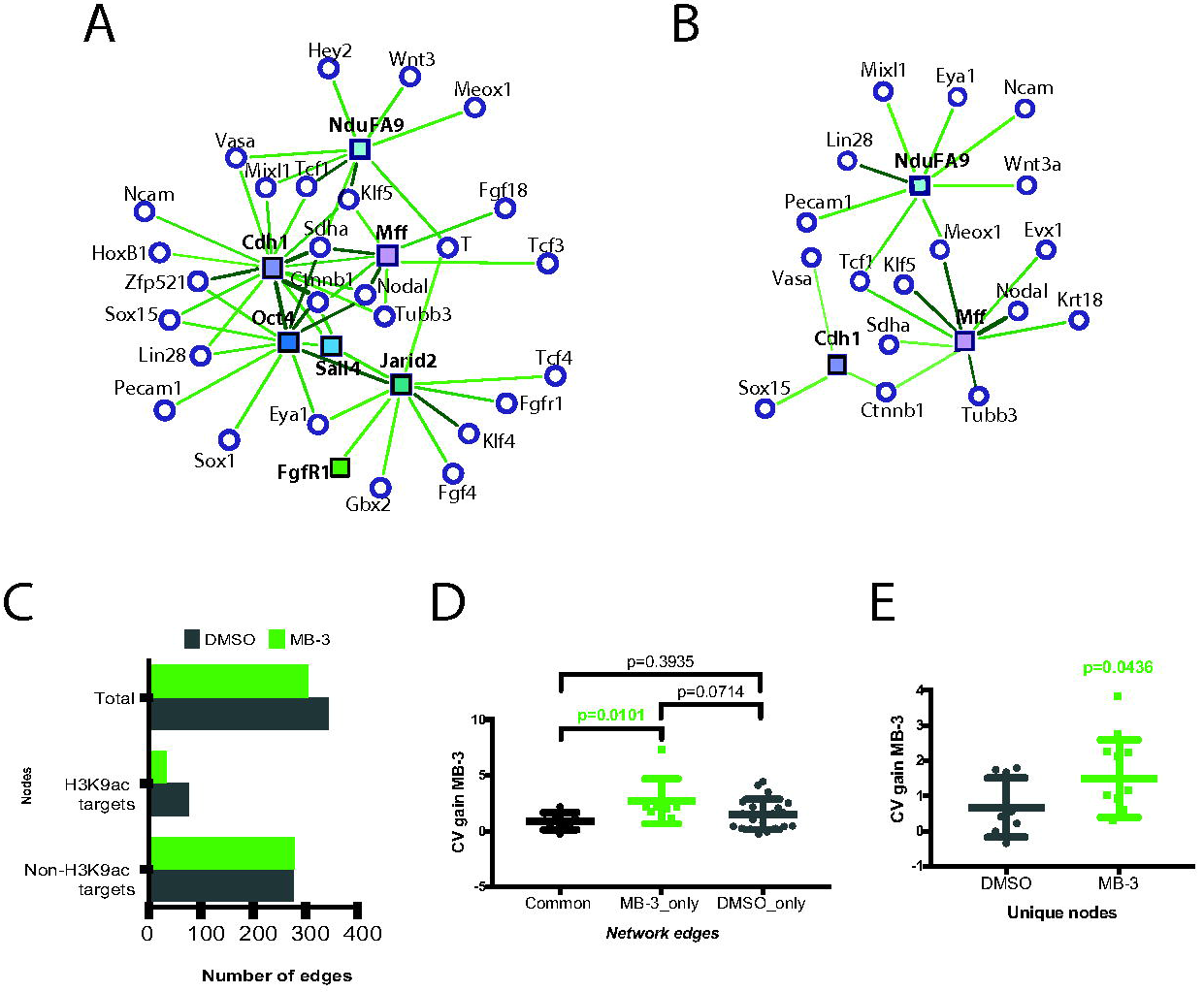
Network analysis of MB-3 treated cells shows destabilisation following Kat2a inhibition. **A-B**. Association network diagrams of Kat2a-dependent H3K9ac target genes upon DMSO (A) and MB-3 treatment (B). Square nodes, Kat2a-dependent H3K9ac target genes; Edge colour, strength of correlation (darker green = stronger correlation); White nodes, correlated genes. **C**. Enumeration of total network edges centred on Kat2a-dependent H3K9ac target and non-target genes upon treatment of TNGA ES cells in 2i with either DMSO or MB-3. **D**. CV gain [(CV_mb-3_-CV_DMSO_)/CV_DMSO_] for DMSO-unique, MB-3 unique and common edges in the remodelled networks centred on Kat2a-dependent H3K9ac targets (shown in A and B). **E**. CV gain in network nodes exclusively present in DMSO or MB-3 treatment of TNGA ES cells in 2i. All analyses used Paired Student’s t-test.

The nature of the genes unique to each network underlies the phenotypes observed. In addition to *Pou5f1*, *Sall4* and *Jarid2*, the control network uniquely also includes Fgf signalling elements, recently reported to be regulated by Kat2a [27], and *Id3*, a BMP target [21]. Amongst the nodes specific to the MB-3 network are endodermal (*Afp*), mesodermal (*Chrd, Wnt3a*) and neuroectodermal (*Dmbx1*, *Evx1*) differentiation genes. Interestingly, the MB-3 network also uniquely includes *Nanog*, whose presence in the network may reflect capture of the proposed early transition state into commitment (Fig. 5 and Supplementary Fig. 3). Overall, our data support the notion that MB-3-mediated loss of H3K9ac at target loci critically reconfigures the architecture of the gene regulatory networks sustaining pluripotency, through a direct effect on variability of gene expression.

## DISCUSSION

We have shown that changes in the activity of the histone acetyl-transferase Kat2a deplete a significant proportion of H3K9 acetylation sites at promoter locations and alter gene expression heterogeneity in mouse ES cells. Affected loci include key components of the pluripotency network, as well as general metabolic categories, which suggests that this mode of action may be extended to other cell types and developmental contexts. We have estimated heterogeneity on the basis of expression level CV and shown that Kat2a-inhibited cultures exhibited higher CVs than controls, particularly when probed from naïve (2i) pluripotency conditions.

Functionally, this translated into a curtailed ability to establish naïve pluripotency and an increased propensity to initiate ME differentiation, best explained by an increased probability to irreversibly exit the pluripotent state, without accompanying changes in cell cycle or apoptosis. Our recent report that loss of KAT2A can promote differentiation of human leukaemic cells [20] is compatible with a generalisation of the principles established in this work.

Notably, the radical changes observed upon MB-3 treatment occur with minimal perturbation of average gene expression levels, sustaining the notion that variability of gene expression can, in itself, alter the probability of state transitions of individual cells. This is compatible with our previous observations in the haematopoietic system that lineage specification can be consequent to distinct individual molecular events, which are stochastic in nature and independently drive re-organisation of transcriptional programmes [9]. Indeed, in mouse ES cells treated with MB-3 and sampled from a newly-established naïve pluripotency state, transcriptional heterogeneity resulted in a dramatic reconfiguration of GRNs, with destabilisation of *Pou5f1/Oct4* connectivity, as well as of other transcriptional and signalling regulators of pluripotency, including transcriptional and cell-cell adhesion components of the Wnt pathway. Furthermore, GRN remodelling observed upon Kat2a inhibition included novel associations suggestive of differentiation, but also concomitant nucleation of the networks by *Nanog*, likely denoting the capture of an early and normally transient state in which differentiation programmes are primed in the context of a subsisting pluripotency network. Our experimental and modelling data lends support to this interpretation.

The underlying transcriptional variability that accompanies capture of the mid-Nanog transition state is reminiscent of other transition events associated with enhanced heterogeneity [2, 3, 5, 7]. Recently, it was suggested that heterogeneity of expression levels occurs primarily in genes which decrease their expression at fate decisions [34], but that this heterogeneity resolves on commitment to a stable, differentiated state, requiring a dynamic means of transcriptional regulation. In line with this view, Ahrends et al. [1] have modelled the contribution of transcriptional noise during commitment and progression and adipocyte differentiation and found that while low levels of noise ensure lineage commitment, noise must be limited for differentiation to progress. Lysine acetylation is a highly dynamic post-translational modification, and selective perturbation of H3K9 acetylation constitutes an interesting regulatory paradigm, as it associates with maintenance, but not with initiation, of transcriptional activity [35]. As such, reduction in H3K9 acetylation at specific loci is likely to destabilise, but not abolish, the transcriptional processes, and result in stochastic gene expression, on which other more stable epigenetic marks can act. At a molecular level, our results mirror the effects of the yeast Kat2a homologue Gcn5 on transcriptional noise [18, 32], and thus suggest that in eukaryotes, H3K9 acetylation may play a central role in the control of heterogeneity. Our observations also suggest that dynamic control of transcriptional noise may be key to efficient state transitions.

Early reports on Kat2a null mouse ES cells did not indicate a requirement for maintenance of pluripotent cultures [36]. However, they did denote an acceleration of differentiation of embryoid bodies, and reduced contribution to chimaeras, both of which are compatible with our observed destabilisation of pluripotency and anticipation of early ME differentiation, as well as with the proposition that persistent high noise levels prevent terminal execution of lineage programmes. Indeed, Kat2a-null embryoid bodies, particularly in a conditional knockout framework, constitute a good system in which to test the contribution of gene expression heterogeneity at different stages of differentiation progression, and in different lineages. Recent detailed analysis of the molecular pathways underlying the Kat2a null defects in embryoid bodies [27] identified an association with general metabolic pathways, which we also see in our work, and a specific defect in Fgf signalling. Although Fgf signalling is actively repressed in our experimental context, it is noteworthy that Fgfr1 is a direct target of Kat2a-mediated H3K9 acetylation, and that Fgf4 and Fgfr1 exhibit reduced network connectivity and increased variability of gene expression upon MB-3 treatment.

Finally, it should be noted that while we validated the cellular consequences of Kat2a inhibition in different mouse ES cells, and with gene expression knockdown, we conducted the molecular analyses in TNGA cells, which allowed for the direct exclusion of committed cells and the verification of increased culture heterogeneity by a simple measure. TNGA are heterozygous for Nanog, and consequently partially defective for the auto-regulatory loop that maintains Nanog expression [16] and re-enforces pluripotency. The loss or inhibition of Kat2a in these cells may be sufficient to affect the metastability of the primed pluripotency state and enhance heterogeneity. The reproducibility of cellular events in non-TNGA cells suggests a more universal mechanism of noise enhancement, but this may need to be experimentally validated in other ES cell lines, and distinct cell types. Along the same line, it will be interesting to test whether other epigenetic regulators suggested to regulate transcriptional noise in yeast [18] have a similar impact on exit from pluripotency, or if sequence-editing of locus-specific regulatory elements targeted by H3K9 acetylation promotes exit from pluripotency in wild-type mouse ES cells. The on-going expansion of CRISPR-based tools for gene and epigenetic editing [37] should assist in these experiments and provide an invaluable resource for dissection of cell state transitions in multiple stem cell systems and during development.

## ACKNOWLEDGEMENTS

We are grateful to Alfonso Martinez Arias for critical reading of the manuscript. We thank staff of Cambridge NIHR BRC Cell Phenotyping Hub, CIMR Flow Cytometry Core Facility and Department of Pathology of the University of Cambridge for expert assistance with cell sorting. Library preparation for next-generation sequencing was performed in the Cambridge Stem Cell Institute Genomics Core facility, with sequencing performed in the Genomics Core Unit of the CRUK Cambridge Institute. NM was funded by a BBSRC DTP Studentship, and SE is supported by a Cambridge Trust Studentship. KR was funded by an Isaac Newton Fellowship to CP and through an INT/WT-ISSF/Univ Cambridge Joint Research Grant to CP. M.F. work is supported by the British Heart Foundation (BHF) Cambridge Centre of Excellence (RE/13/6/30180); D.S. work has been supported by an Isaac Newton fellowship to M.F. CP is funded by a Kay Kendall Leukaemia Fund Intermediate Fellowship (KKL888) and a Leuka John Goldman Fellowship for Future Science (2017).

## DISCLOSURE OF POTENTIAL CONFLICTS OF INTEREST

The authors have no conflicts of interest to disclose.

---

**Supplementary Figure 1.**
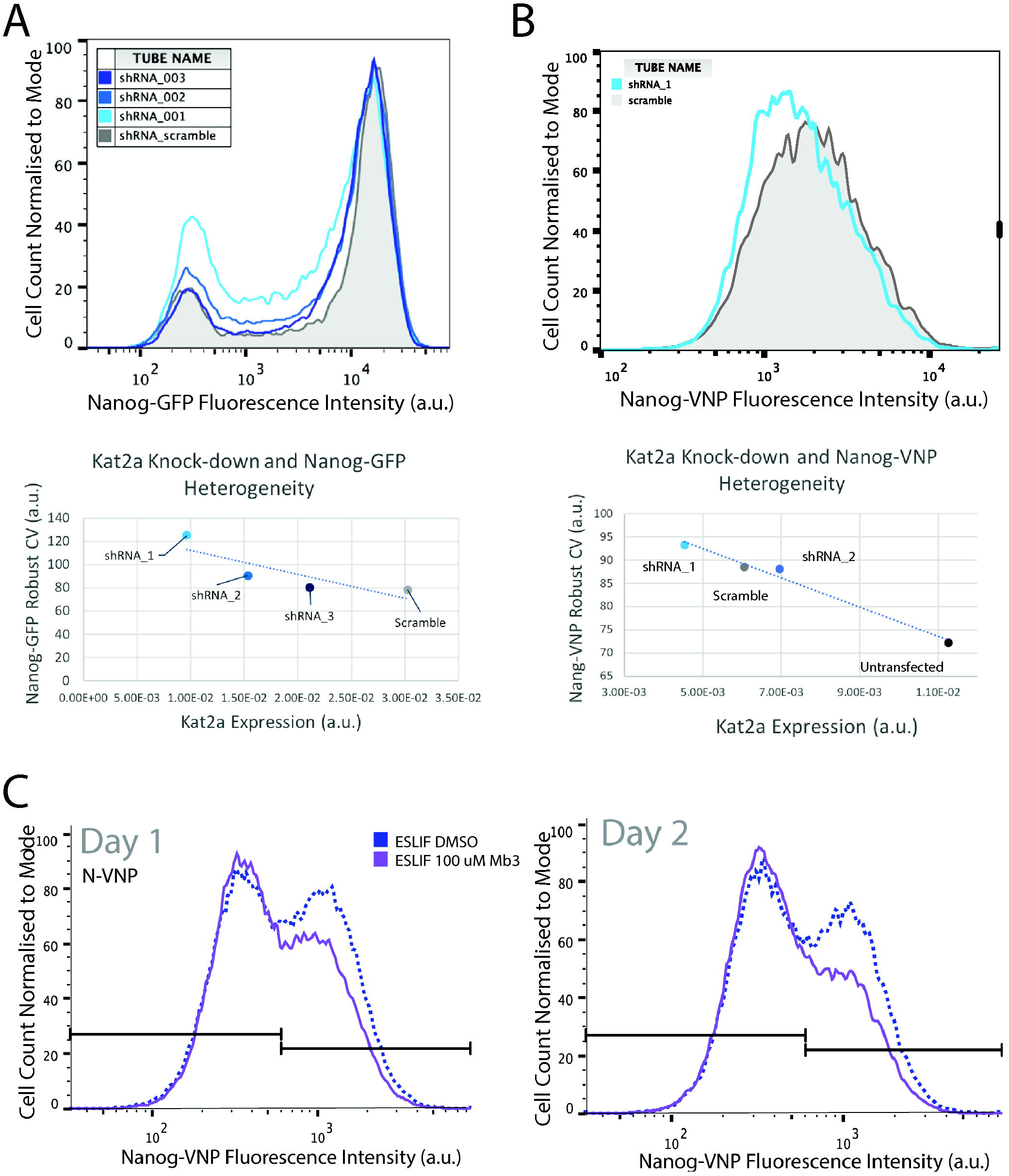

**Supplementary Figure 2.**
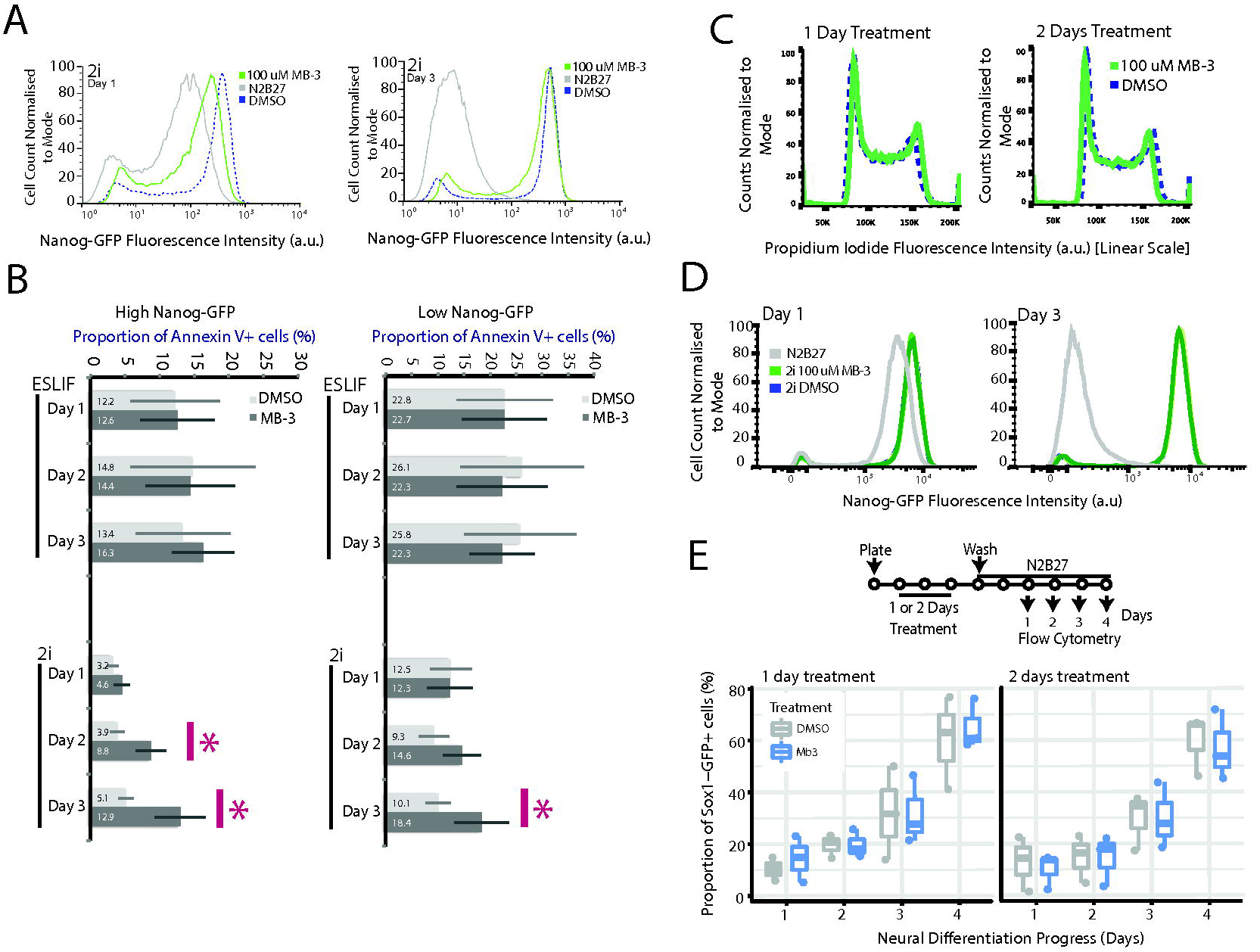

**Supplementary Figure 3.**
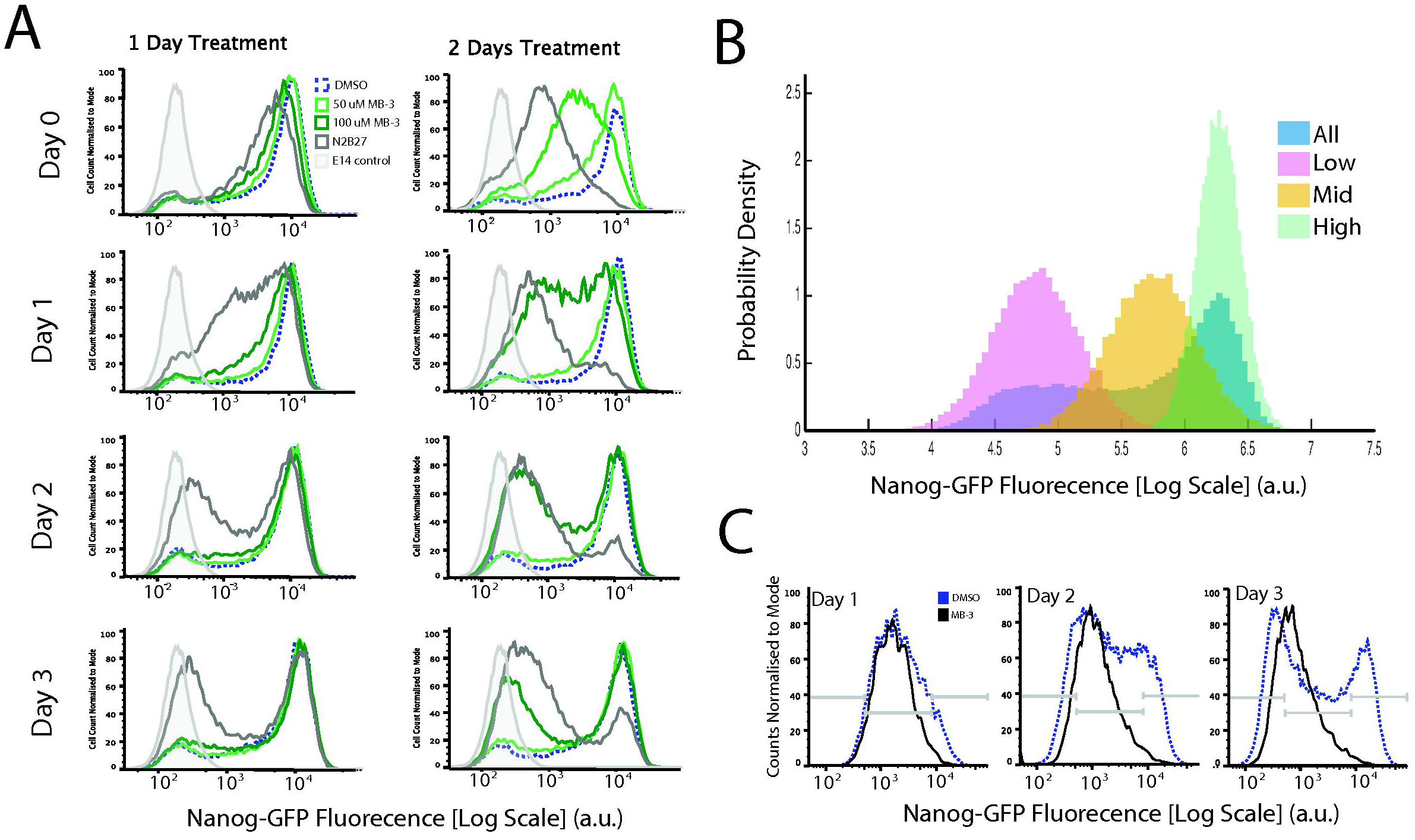

**Supplementary Figure 4.**
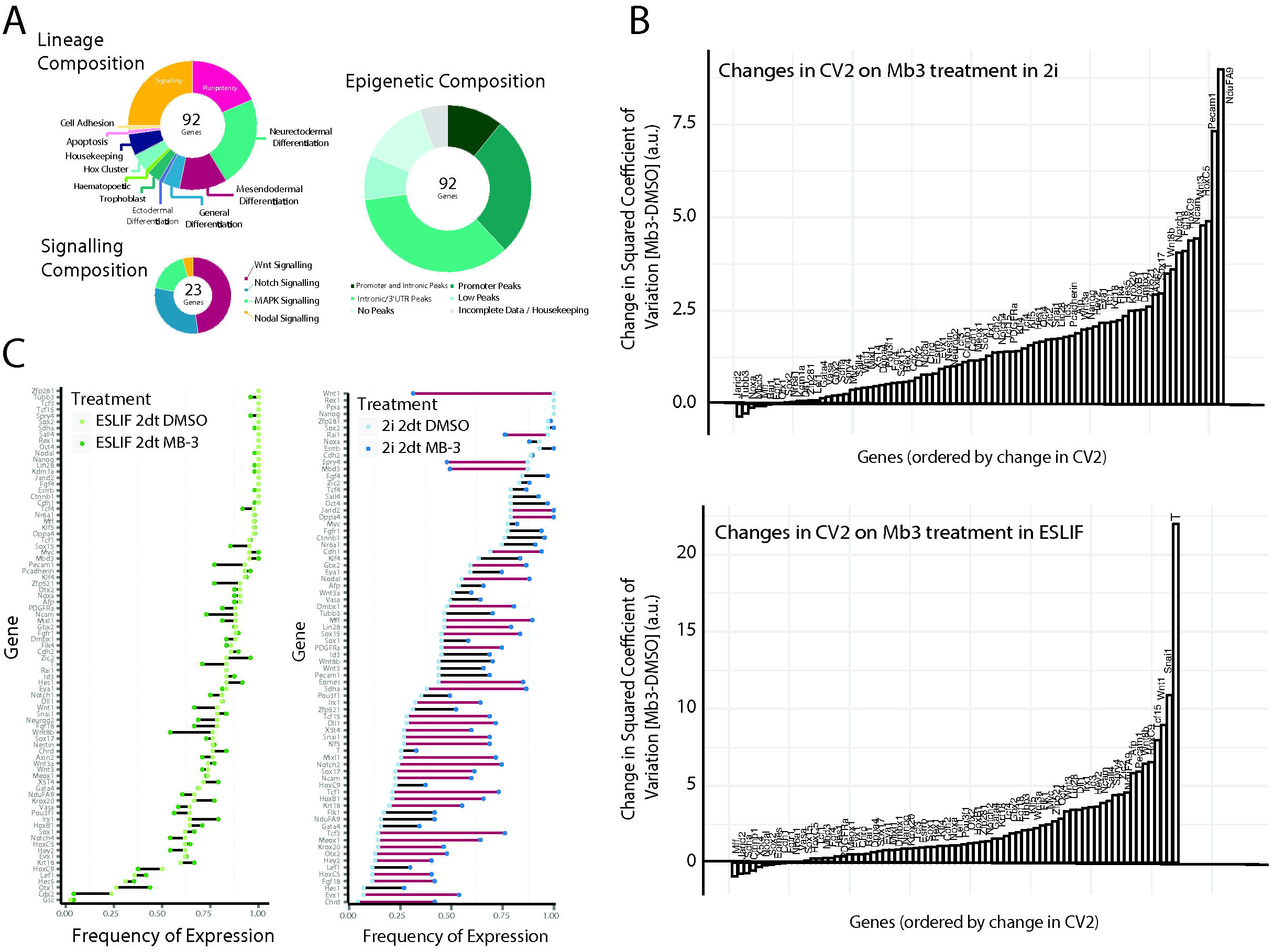

